# Mechanisms of adaptive interlimb coordination to sudden ground loss: a neuromusculoskeletal modeling study

**DOI:** 10.1101/2025.11.11.687930

**Authors:** Kota Shinohara, Yuichi Ambe, Yongi Kim, Fu Mano, Shravan Tata Ramalingasetty, Andrew B. Lockhart, Sergey N. Markin, Jessica Ausborn, Ilya A. Rybak, Simon M. Danner, Shinya Aoi

## Abstract

Mammals exhibit robust walking across diverse environments, a capability largely attributed to central pattern generators (CPGs) in the spinal cord. Afferent feedback modulates CPG output and plays a critical role in adaptive locomotion, yet its specific contributions remain poorly understood. To investigate this, we used a neuromusculoskeletal model to simulate hindlimb locomotion in spinalized cats encountering a hole and experiencing a sudden loss of ground support, as described in prior experimental studies. The model couples a trunk-and-hindlimb musculoskeletal system to a pair of two-level, half-center CPGs—one for each hindlimb. The model reproduced the observed adaptive interlimb coordination that allows cats to maintain walking after the sudden loss of ground support. Notably, the adaptive response emerged without re-optimizing parameters, which were tuned for steady walking in an environment without holes. Nullcline analysis based on dynamical systems theory revealed that afferent feedback mechanisms controlling the transitions between fast and slow neuronal dynamics facilitated adaptive interlimb coordination. These findings provide mechanistic insight into how spinal feedback circuits support robust locomotion through dynamic interactions between the nervous system, the musculoskeletal system, and the environment.

## 1. Introduction

Mammals skillfully control their complex musculoskeletal systems, enabling robust locomotion across diverse environments. This ability relies on spinal neural circuits that generate rhythmic motor patterns in response to descending commands and adapt them through afferent feedback integration. These circuits are commonly referred to as central pattern generators (CPGs) [18, 22, 46, 40, 47].

The CPGs are capable of autonomous activity even in the absence of sensory feedback. When continuous electrical stimulation is applied to the mesencephalic locomotor region in immobilized decerebrate cats, alternating activity of flexor and extensor motoneurons (MNs) is observed, similar to that during walking in intact cats, and is called fictive locomotion [46]. Fictive locomotor preparations, combined with sensory nerve stimulation have been used to investigate the adaptive regulation of locomotor patterns by somatosensory afferent feedback [23, 43, 37, 55]. To elucidate the regulation mechanism mathematically, a two-level half-center CPG model comprising a rhythm generator (RG) layer and a pattern formation (PF) layer has been proposed [48, 49]. This model was later refined to study speed-dependent gait regulation in mammals [3, 5, 6, 7, 32, 38, 50, 51, 52, 64].

Mammalian locomotion is a complex behavior resulting from dynamic interactions among the nervous system, musculoskeletal system, and environment. Understanding adaptive locomotion requires studying these systems in concert, including the role of sensory feedback and biomechanical context. Decerebrate and spinal cat models have been instrumental in revealing how afferent feedback from body dynamics modulates neural control. In addition, perturbations such as unexpected holes [25], obstacles [65], and external forces [31], as well as split-belt treadmills [33, 15, 16, 17, 30, 62, 63] have helped characterize the principles of adaptive motor control of locomotion. These studies have shown that mammals exhibit robust adaptive responses to environmental perturbations. However, the mechanisms of adaptation through the dynamic interactions between the nervous system, the musculoskeletal system, and the environment remain poorly understood.

Experimental studies in spinal cats have shown that a sudden loss of ground support—such as stepping into a hole during treadmill walking—elicits a rapid, adaptive motor response [25]. When the foot entered the hole during locomotion and the hindlimb becomes hyperextended, flexor muscle activity is initiated early to lift the foot out of the hole and resume walking [25]. Critically, this recovery also requires sustained extensor activity in the contralateral hindlimb to support body weight. As a result, two flexions occur on the affected side-one before entering the hole and one during recovery-while the opposite limb maintains extension. This disruption of the typical left-right alternation underscores the importance of interlimb coordination in adaptive locomotion.

Despite these insights, the dynamic mechanisms that enable such coordinated responses remain poorly understood. We previously investigated the mechanism underlying the rapid flexion of the ipsilateral hindlimb using a neuromusculoskeletal model of a single hindlimb [28]. In the present study, we built on this framework by developing a bilateral neuromusculoskeletal model that incorporates structured afferent feedback and physiologically-inspired spinal circuitry controlling both hindlimbs and their coordination. Using simulation and dynamical systems analysis, we examined how spinal circuits leverage sensory input to flexibly reorganize motor output following perturbation. Our approach provides mechanistic insight into how spinal feedback circuits support stable locomotion through context-dependent modulation of interlimb coordination.

## 2. Model

### 2.1. Musculoskeletal model

We have modified the cat neuromusculoskeletal model from our previous study [28], as shown in Fig. 1. The skeletal model is two-dimensional and consists of seven rigid links representing a trunk and both hindlimbs. Each hindlimb consists of a thigh, crus, and foot connected by hip, knee, and ankle joints. The forelimbs are fixed to the trunk and the tips are constrained to a stationary platform 2 cm above the treadmill through horizontal viscoelastic elements. The model walks on the treadmill with a belt speed of 20 cm/s. When the trunk is parallel to the belt and the thigh, crus, and foot are in a straight line and perpendicular to the trunk, the hip angle is 135° and the knee and ankle angles are both 180°. The joint angles increase as the joints are extending. The contact between the hindlimb tip and treadmill was modeled using viscoelastic elements. We derived the equations of motion using Lagrangian equations with the same physical parameters for the skeletal model as those in [13] and solved the equations numerically using the fourth-order Runge-Kutta method with a time step of 0.04 ms.

**Figure 1.**
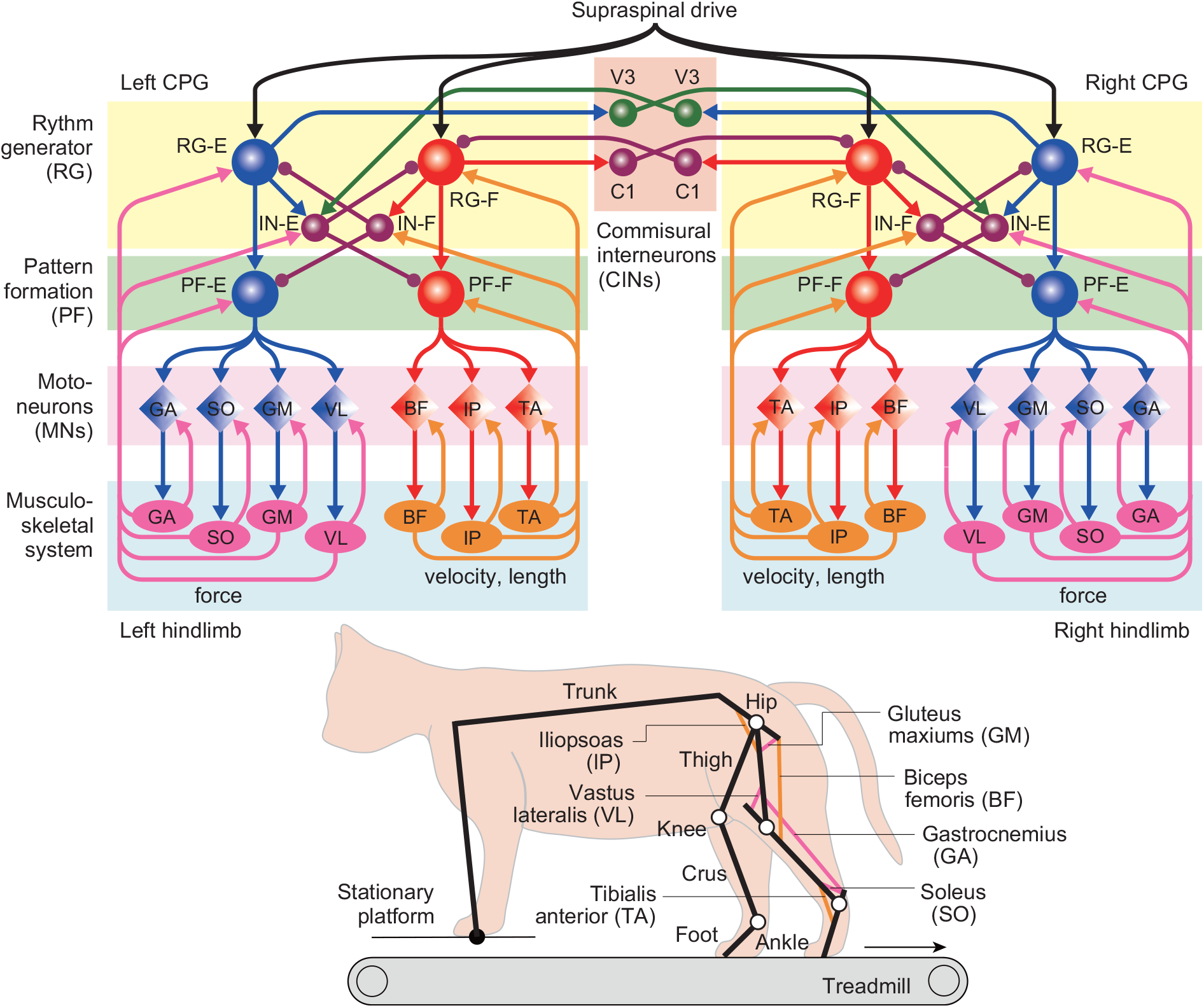
Neuromusculoskeletal model of a cat walking on a treadmill using only the hindlimbs, with the forelimbs stabilized on a stationary platform.

We used seven Hill-type muscles for each hindlimb, where five muscles are uni-articular: hip flexor (iliopsoas, IP), hip extensor (gluteus maximus, GM), knee extensor (vastus lateralis, VL), ankle flexor (tibialis anterior, TA), ankle extensor (soleus, SO), and two muscles are bi-articular: hip extensor/knee flexor (biceps femoris, BF) and knee flexor/ankle extensor (gastrocnemius, GA). We assumed that the moment arms of the muscles are constant. We used the same muscle model to generate tension through contractile and passive elements as in [13]. Specifically, the muscle force 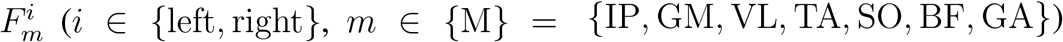 is given by

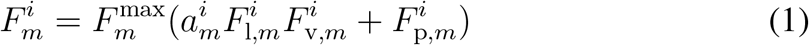

where 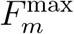 is the maximum force, 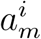 is the muscle activation 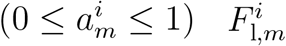, is the force-length relationship, 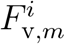 is the force-velocity relationship, and 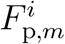 is the passive component. The muscle lengths were normalized by 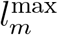, which was set so that the uni-articular muscles had a length of 85% of 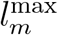 and bi-articular muscles were at 75% at a neutral posture with the hip joint at 60°, the knee joint at 90°, and the ankle joint at 100°. In addition, 2° of joint motion corresponded to 1% of muscle length change, except for the GA muscle, where 1.5° at the ankle joint or 4.5° at the knee joint was required. The muscle velocities were normalized by 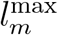.

The muscle activation 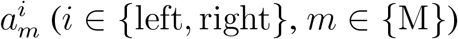 is determined through a low-pass filter [61] given by

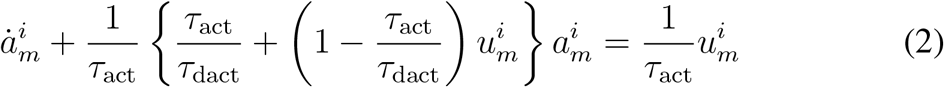

where *τ*_act_ (= 20 ms) and *τ*_dact_ (= 32 ms) are activation and deactivation time constants, respectively, and 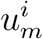 is the motor command determined from the corresponding motoneuron activity of the motor control model.

### 2.2. Motor control model

It has been suggested that the locomotor behavior is controlled by the CPG, which consists of hierarchical networks with rhythm generator (RG) and pattern formation (PF) networks [48, 49]. The RG network produces the rhythmic activity and the PF network produces the spatiotemporal patterns of motor commands. We developed a motor control model based on our previous two-level half-center CPG model for a single cat hindlimb, comprising RG and PF networks [28]. Specifically, we integrated two CPG models that control the left and right hindlimbs, as shown in Fig. 1. The RG of each CPG model has two neuronal populations that represent the flexor and extensor centers (RG-F and RG-E), which receive a supraspinal drive, and two populations of inhibitory interneurons (IN-F and IN-E), which provide mutual inhibition between the RG-F and RG-E. The RGs of the left and right CPG models are connected through commissural interneurons (CINs) based on [38]. The CINs have the population of C1 neurons, which mediate the inhibition of the contralateral RG-F by the ipsilateral RG-F, and the population of V3 neurons, which mediate the inhibition of the contralateral RG-F by the ipsilateral RG-E through the contralateral IN-E. Like the RG, the PF also has two neuronal populations that represent the flexor and extensor centers (PF-F and PFE), which receive the excitatory input from the RG neurons and inhibitory input from the INs. Each CPG model has seven populations of motoneurons (MNs) that provide the motor command for the corresponding muscle (MN-*m, m* ∈ *{*M*}*). The MNs of flexor muscles MN-IP, MN-TA, and MN-BF receive excitatory input from the PF-F, while those of extensor muscles MN-GM, MN-VL, MN-SO, and MN-GA receive the excitatory input from the PF-E. Each CPG model receives afferent feedback from the ipsilateral muscles.

We used the activity-based (non-spiking) neuron model [14, 34, 38, 5, 6] for each population. The state of each neuron is characterized by the membrane potential 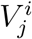 (*i* ∈ *{*left, right*}, j* ∈ *{*RG*}, {*IN*}, {*CIN*}, {*PF*}, {*MN*}*), where *{*RG*}* = *{*RG-F, RG-E*}, {*IN*}* = *{*IN-F, IN-E*}, {*CIN*}* = *{*C1, V3*}, {*PF*}* = *{*PF-F, PF-E*}*, and *{*MN*}* = *{*MN-*m* | *m* ∈ *{*M*}}*. In RG, PF, and MN, a persistent (slowly inactivating) sodium current determines the intrinsic rhythmogenic properties of these neurons. Therefore, the RG, PF, and MN use the variable 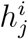 (*i* ∈ *{*left, right*}, j* ∈ *{*RG*}, {*PF*}, {*MN*}*) that defines the slow inactivation of the persistent sodium channel. The RG-F and RG-E produce rhythmic activity. However, when they are uncoupled, the RG-E is in the tonic regime due to the supraspinal drive and produces sustained activity. Rhythmic oscillation of the RG is defined by the RG-F, which provides rhythmic inhibition of the RG-E through the IN-F. The oscillation frequency is determined by the supraspinal drive to the RG-F. When the PF and MN are uncoupled, they do not show rhythmic activity due to the relatively low maximum conductance of the sodium current. They produce rhythmic activity through the excitatory inputs from the RG.

For the state variable, we used 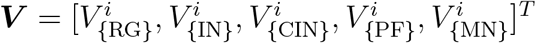 and 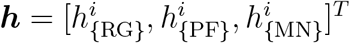. The dynamics of the membrane potential 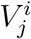 is given by

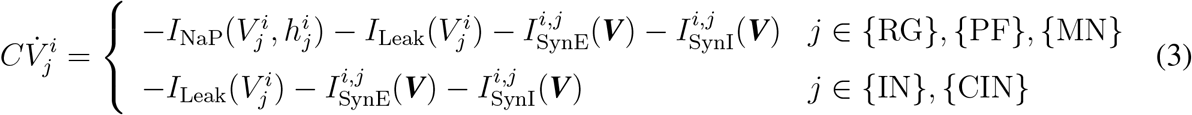

where *C* is the membrane capacitance, *I*_NaP_ is the persistent sodium current, *I*_Leak_ is the leakage current, and 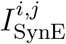 and 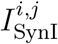are the currents from excitatory and inhibitory synapses, respectively. The ionic current *I*_NaP_ and leakage current *I*_Leak_ are given by

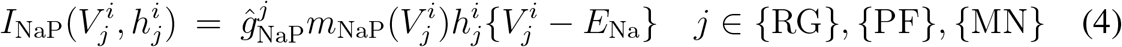

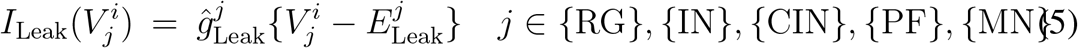

where 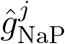 and 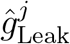 are the maximum conductances, and *E*_Na_ and 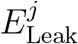 are the reversal potentials. *m*_NaP_ is the activation of the sodium channel of the RG, PF, and MN and is given by

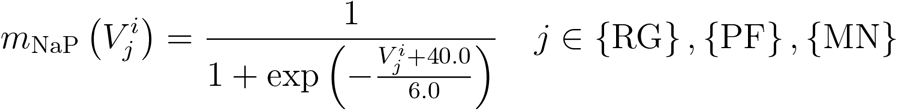

The dynamics of the inactivation of the sodium channel 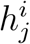 of the RG, PF, and MN is given by

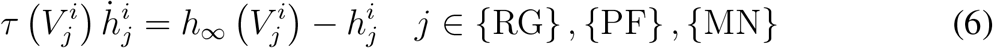

Where

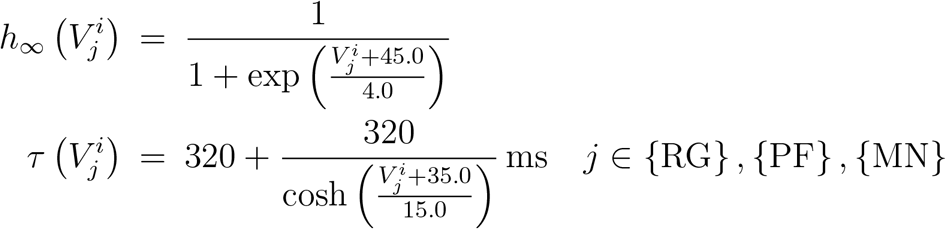

The currents induced by the synapses 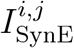 and 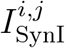 are given by

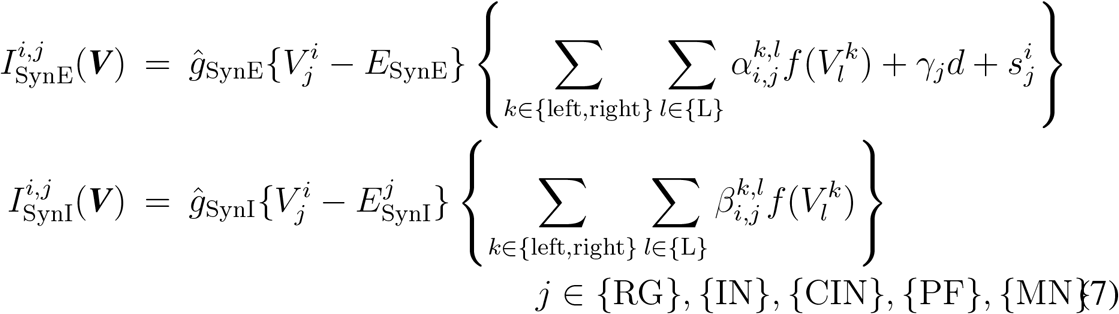

where 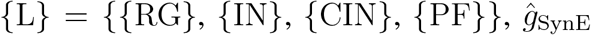 and 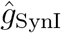 are the maximum conductances, *E*_SynE_ and 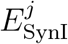 are the reversal potentials, *d* is the tonic drive from the supraspinal region, and 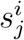 is the afferent feedback from the musculoskeletal model, as described below. 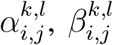, and *γ*_*j*_ are the weight coefficients of the excitatory, inhibitory, and supraspinal connections, respectively, where 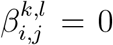 for *j* ∈ *{*CIN*}, {*MN*}* and *γ*_*j*_ = 0 for *j* ∈ *{*IN*}, {*CIN*}, {*PF*}, {*MN*}*. The output function *f* translates *V* into the integrated population activity and is given by

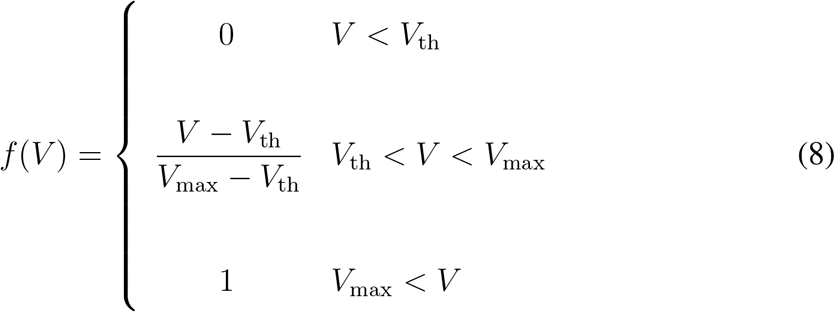

where *V*_th_ and *V*_max_ are the lower and upper threshold potentials, respectively. The motor command 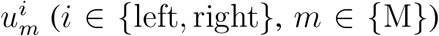 is given by 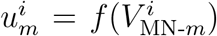. Based on [20, 28, 34], we determined the parameters for the motor control model (see Appendix A) except for 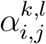 for *j* ∈ *{*MN*}* and *l* ∈ *{*PF*}*, which we determined through optimization, as described below.

The afferent feedback 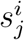 is given by

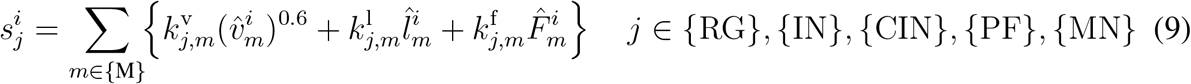

Where

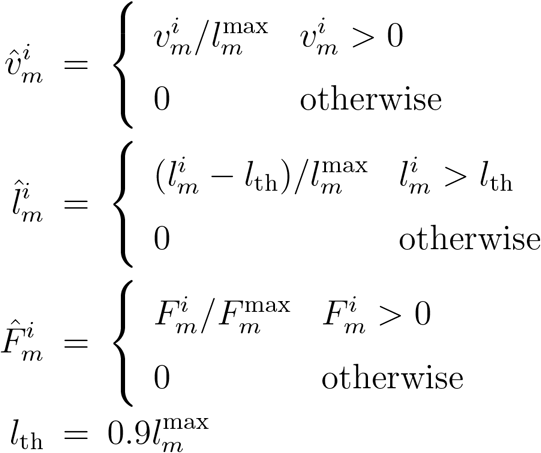

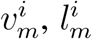, and 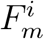 are the velocity, length, and force of muscle *m*, respectively, and 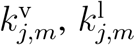, and 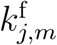 are the corresponding weight coefficients. The first, second, and third terms of (9) represent the velocity, length, and force feedback from muscle *m*, respectively. In particular, we used the velocity and length feedback only for the flexor muscles and the force feedback only for the extensor muscles to focus on the adaptation to the loss of ground support during walking. Specifically, the coefficients 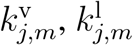, and 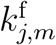 for the flexor muscles (*m* ∈ *{*IP, TA, BF*}*) are given by

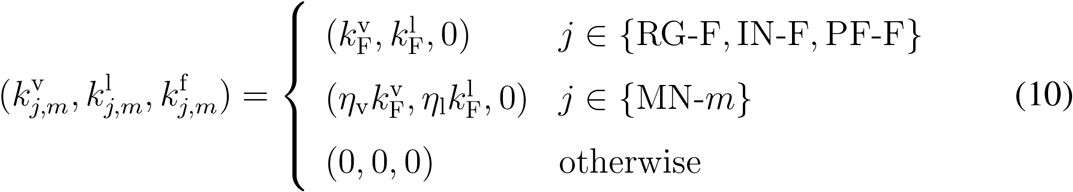

Those for the extensor muscles (*m* ∈ *{*GM, VL, SO, GA*}*) are given by

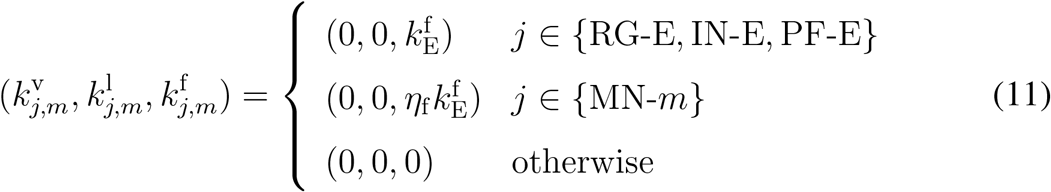

We determined 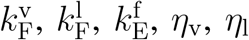, and *η*_f_ through the optimization as described below.

### 2.3. Parameter determination by optimization

We determined the parameters of the motor control model by optimization. To investigate the general role of sensory feedback in the response to perturbations, we used a walking environment without any holes, but with randomized stepping surface, where the ground level of the treadmill belt of each hindlimb changes randomly among −2, 0, and 2 cm in each step (Fig. 2A).

**Figure 2.**
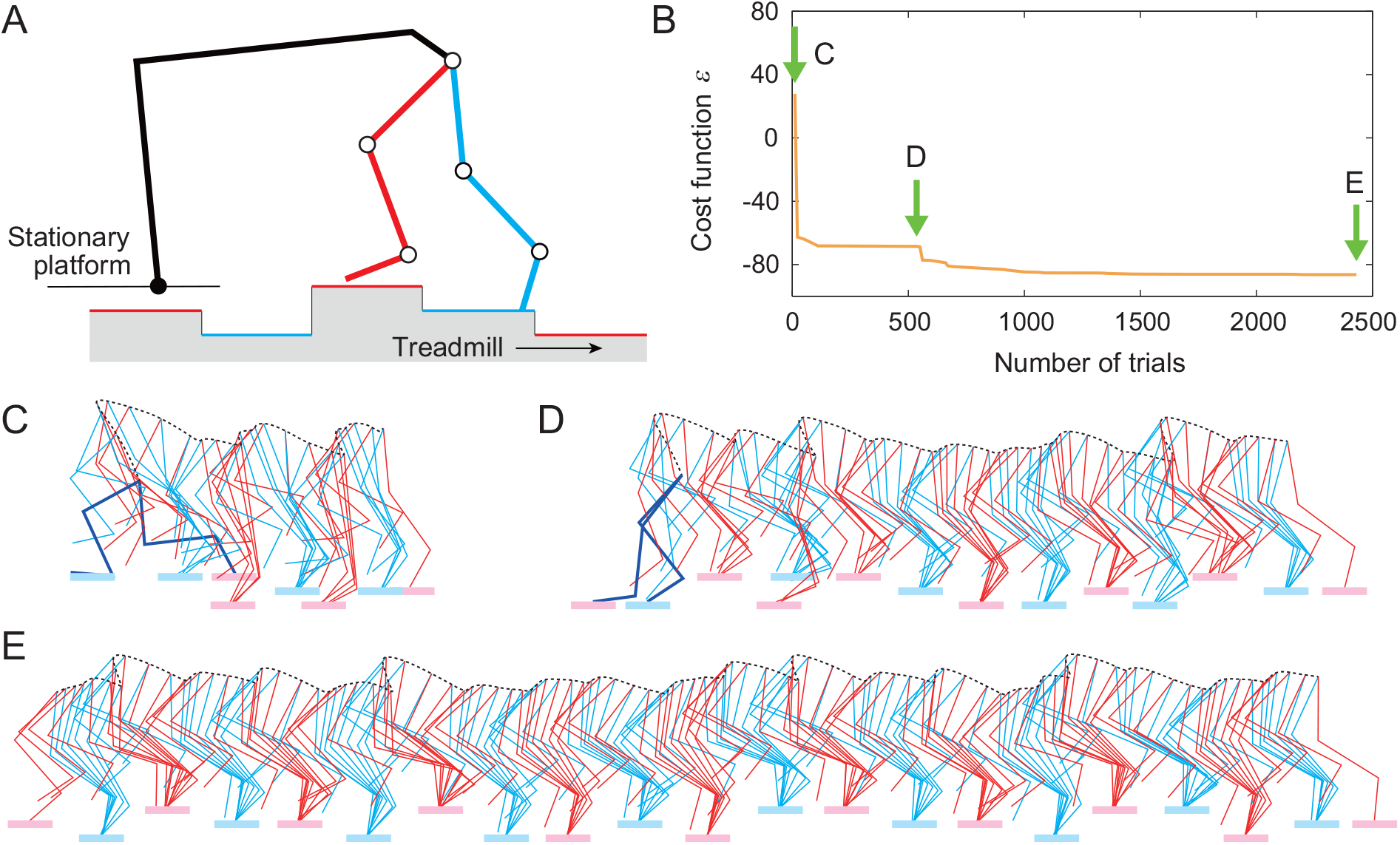
Parameter optimization in an environment with step-to-step random variations in belt height. A. Schematic diagram. B. Cost function for the trials during the optimization process. Stick diagrams at the initial (C), intermediate (D), and final (E) stages of optimization (see Movies 1-3).

We determined the parameters of the motor control model by optimization using a walking environment, where the ground level of the treadmill belt of each hindlimb changes randomly among −2, 0, and 2 cm in each step (Fig. 2A). Specifically, we determined 13 parameters (seven parameters 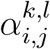 for *j* ∈ *{*MN*}* and *l* ∈ *{*PF*}* to determine the connections from the PFs to the MNs and six parameters 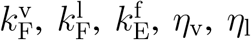, and *η*_f_ to determine the gains of the afferent feedback) using the covariance matrix adaptation evolution strategy (CMA-ES) [24] as the optimization method. To walk for a long time with low muscle activity in the environment, we minimized the following cost function:

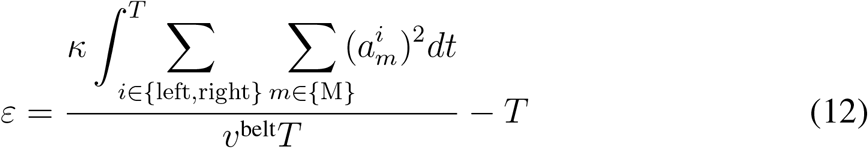

where *κ* = 20, *v*^belt^ (= 20 cm/s) is the belt speed, and *T* is the amount of time the model is able to walk without falling (maximum is 100 s), where we defined falling as when the hip height is below 5 cm and when the foot never touches the belt within one step.

Figure 2B shows the values of the cost function for the trials in the optimization process. The model is initially unable to walk, as shown in Fig. 2C (see Movie 1). However, as optimization progresses, the model can walk for longer durations (Fig. 2D, see Movie 2). Finally, it becomes able to walk stably without falling even in the environment with randomized steps (Fig. 2E, see Movie 3). Appendix B shows the parameters determined in this optimization.

## 3. Simulation results

### Reproduction of flat-surface treadmill walking

We performed walking simulations in which the belt level did not change, using the parameters determined by the above optimization (see Movie 4). Figures 3A and B show the membrane potentials of the RG, IN, CIN, PF, and MN, and the velocity, length, and force feedback from the flexor and extensor muscles. The rhythmic activity of the RG was transmitted to the MN via the PF. Figures 3C and D show the joint angles and muscle activities, respectively, compared with the measured data in cats [44]. Although the left and right hindlimbs show slightly different patterns, the simulated joint angles and muscle activities show similar patterns to the measured data, despite the fact that no measured data were used in the parameter optimization at any stage.

**Figure 3.**
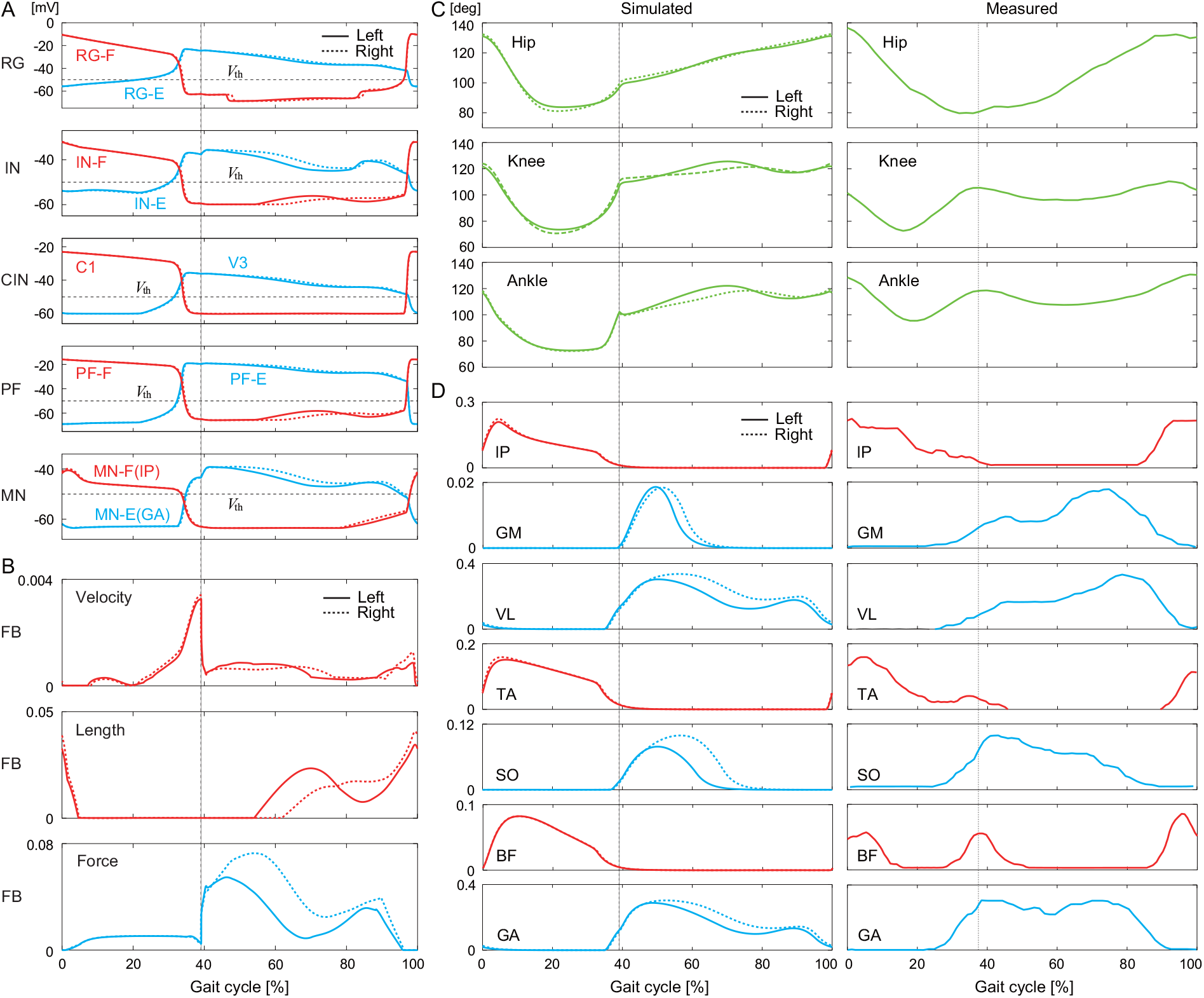
Simulation results for flat-surface treadmill walking using the parameters determined by optimization (see Movie 4). A. Membrane potentials of CPG neurons and B. afferent feedback from flexor and extensor muscles. C. Joint angles and D. muscle activities compared with measured data in cats (adapted from [44]). Liftoffs are represented by 0% and 100% in the gait cycle. Vertical lines indicate touchdowns.

### 3.2. Response to loss of ground support during walking

We used our model with the parameters determined by the above optimization to perform simulation of locomotion when a hole appeared on the left side of the treadmill as in [25]. In our simulation, when a foot entered a hole, it was quickly lifted out of the hole and the model continued to walk on the treadmill without falling (see Movie 5). We compared the simulation results with the measured data in cats [25]. Figure 4A compares the response of the ipsilateral side that fell into the hole. The hip, knee, and ankle joints were first extended and then quickly flexed to lift the foot above the belt for landing. In addition, the onset of the flexor IP muscle activity was advanced. Figure 4B compares the response of the contralateral side. The liftoff position of the foot was further back than during flat-surface walking. In addition, the extensor VL muscle activity was prolonged. These responses on the ipsilateral and contralateral sides were qualitatively similar between the simulation results and the measured cat data.

**Figure 4.**
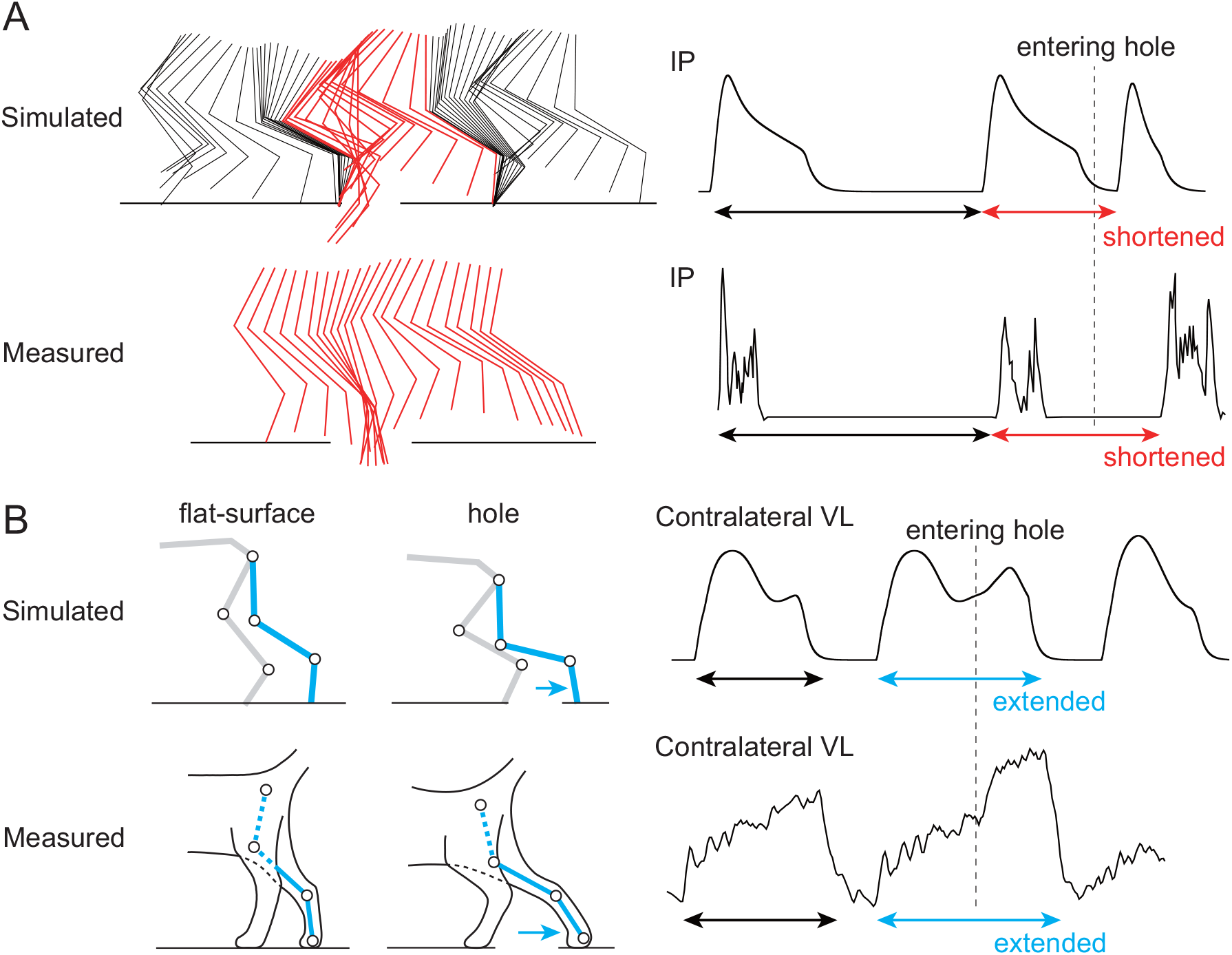
Comparison of response after a foot enters a hole between simulation results (see Movie 5) and cat measured data (adapted from [25]). A. Response on the ipsilateral side that fell into the hole: Stick diagram and IP muscle activity for a few steps before and after entering a hole. The stick diagram of the measured data shows only one step from liftoff to touchdown. B. Response on the contralateral side: Stick diagram and VL muscle activity. The stick diagram compares the posture at liftoff between flat-surface walking and response to entering a hole.

When we used a level surface instead of a randomized stepping surface for the optimization, the model fell over after entering a hole and could not recover from the perturbation. To clarify the role of afferent feedback in responses to the sudden loss of ground support, we performed simulations with afferent feedback removed from the model. Specifically, we tested models lacking velocity feedback from flexor muscles, length feedback from flexor muscles, or force feedback from extensor muscles. We re-optimized the parameters of each model on a randomized stepping surface, as performed above for the model with all feedback intact. We then examined the response of each model when entering a hole. We found that removing any single type of feedback from the model had almost no effect on flat-surface treadmill walking (see Appendix C). However, the models without length or without force feedback fell after entering a hole. Specifically, the model without length feedback could not lift the foot after entering the hole (Fig. 5A, see Movie 6). The model without force feedback exhibited early flexion of the affected hindlimb, but could not support the body weight with the contralateral hindlimb (Fig. 5B, see Movie 7). In contrast, the model without velocity feedback recovered walking after entering the hole in the same way as the model with all feedback (Fig. 5C, see Movie 8).

**Figure 5.**
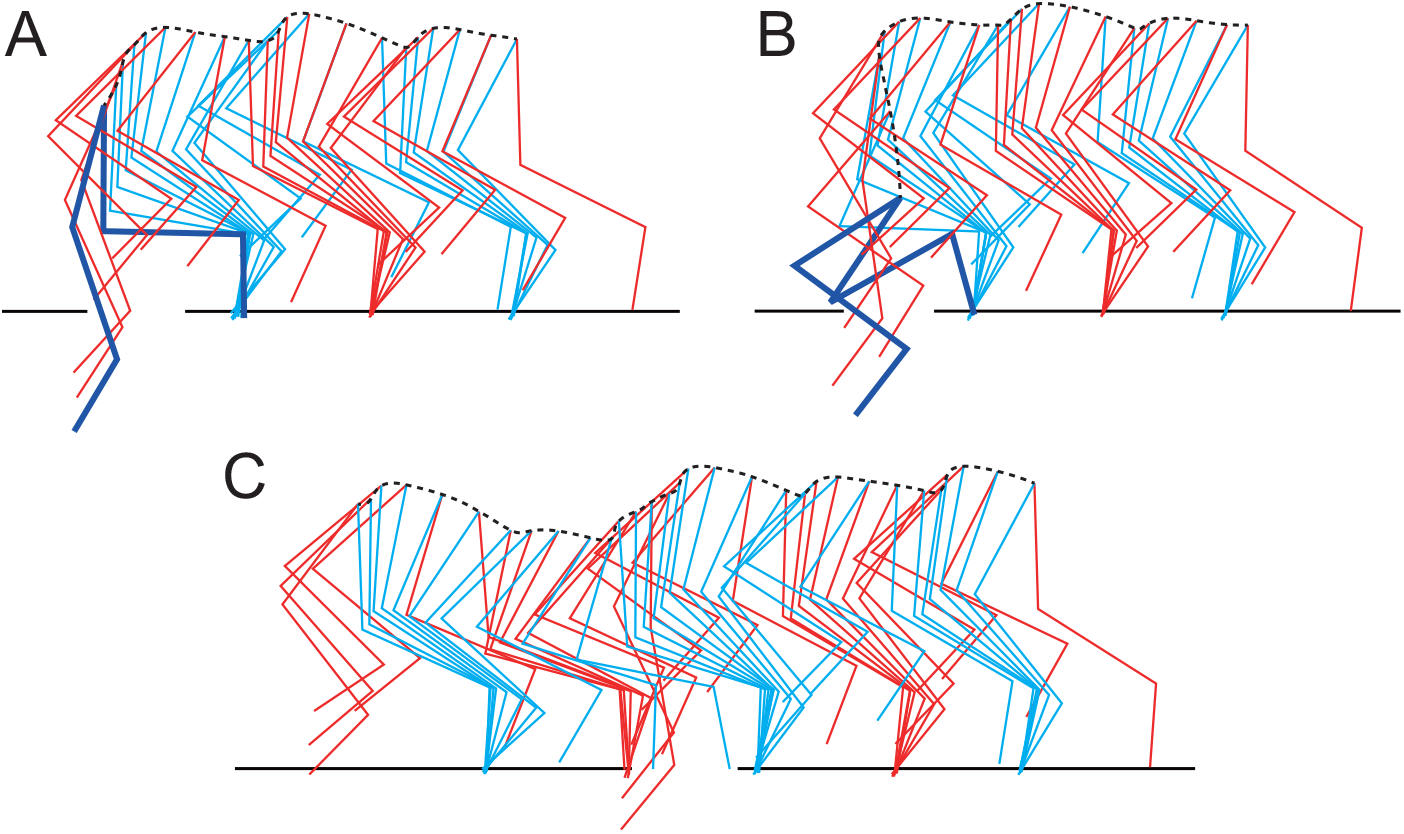
Response after a foot enters a hole for the models without sensory feedback. A. Model without length feedback from the flexor muscles (see Movie 6). The affected hindlimb is not flexed to pull the foot out of the hole. B. Model without force feedback from the extensor muscles (see Movie 7). The contralateral hindlimb cannot support the body weight. C. Model without velocity feedback from the flexor muscles (see Movie 8). This recovers walking after entering the hole.

## 4. Adaptation mechanism

The neuromusculoskeletal simulation showed that the length feedback of the flexor muscles of the left hindlimb (upon entering a hole) and the force feedback of the extensor muscles of the contralateral right hindlimb play an important role in the response to recovery after entering a hole. We, therefore, performed a nullcline analysis to elucidate the mechanism of the CPG responses based on our previous works [20, 28, 54]. Specifically, we investigated the dynamics of the CPG models by focusing on the nullclines of the left RG-F and the right RGE using the simulation results with all afferent feedback during the response in Fig. 4 and compared them with those during flat-surface walking in Fig. 3.

### 4.1. Calculation of nullcline

The nullcline is a set of points at which the derivative of a differential equation is zero and reflects the structure of the solution of the differential equation. Since we found that the length feedback of the flexor muscles of the left hindlimb that entered a hole and the force feedback of the extensor muscles of the contralateral right hindlimb play an important role in the response to recovery after entering a hole from neuromusculoskeletal simulations, we focused on the nullclines of the left RG-F and the right RG-E to clarify the mechanism of the CPG responses and investigated them based on our previous works [20, 28, 54].

The state of the CPG model is given by (***V***, ***h***), and the nullclines for the left RG-F and right RG-E are given by

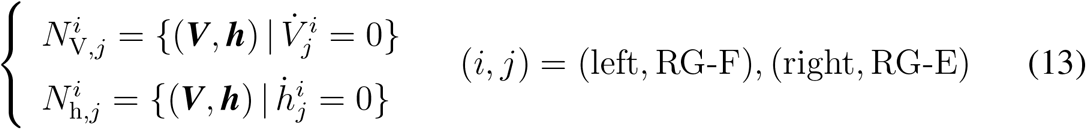

To clarify the dynamics of each left RG-F and right RG-E, only 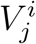 and 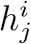for (*i, j*) = (left, RG-F), (right, RG-E) were derived to calculate the nullclines, while the other variables were derived using the variables during walking. Specifically, 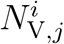 and 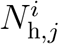 were modified as

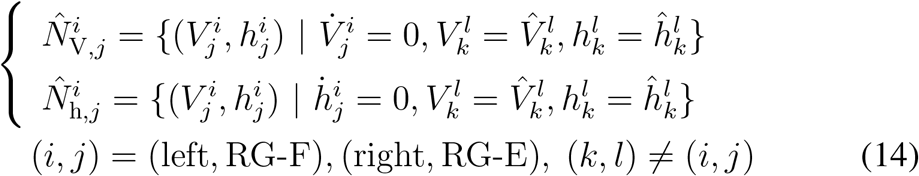

where 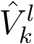and 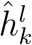 indicate 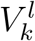 and 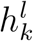 during walking.

While 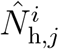 has a sigmoid shape and does not change over time, 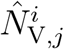has a cubic curve shape and changes over time due to afferent feedback and input from other neurons. Specifically, 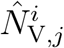 has two different situations with and without two extremes, as shown in Figs. 6A and B. When 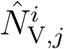 has two extremes (Fig. 6A), the sign of the slope changes and the intersection of 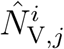 and 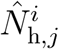 is a saddle. Since the time constant for the dynamics of 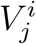 is smaller than that for 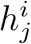, close to where 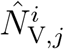 has a positive slope, the state 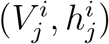 is slowly attracted to the extreme along 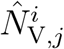 (slow dynamics). Near the extreme, the state jumps to the opposite part of 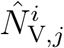 with a positive slope (fast dynamics). When 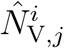 does not have two extremes (Fig. 6B), it changes monotonically and the intersection of 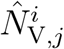 and 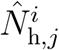 is a stable node. When the state is away from 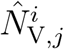, it is quickly attracted to 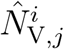 (fast dynamics). Near 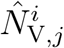, the state is slowly attracted to the stable node along 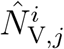 (slow dynamics). Adaptive responses are achieved by switching between fast and slow dynamics depending on the relationship between the state and the nullcline.

**Figure 6.**
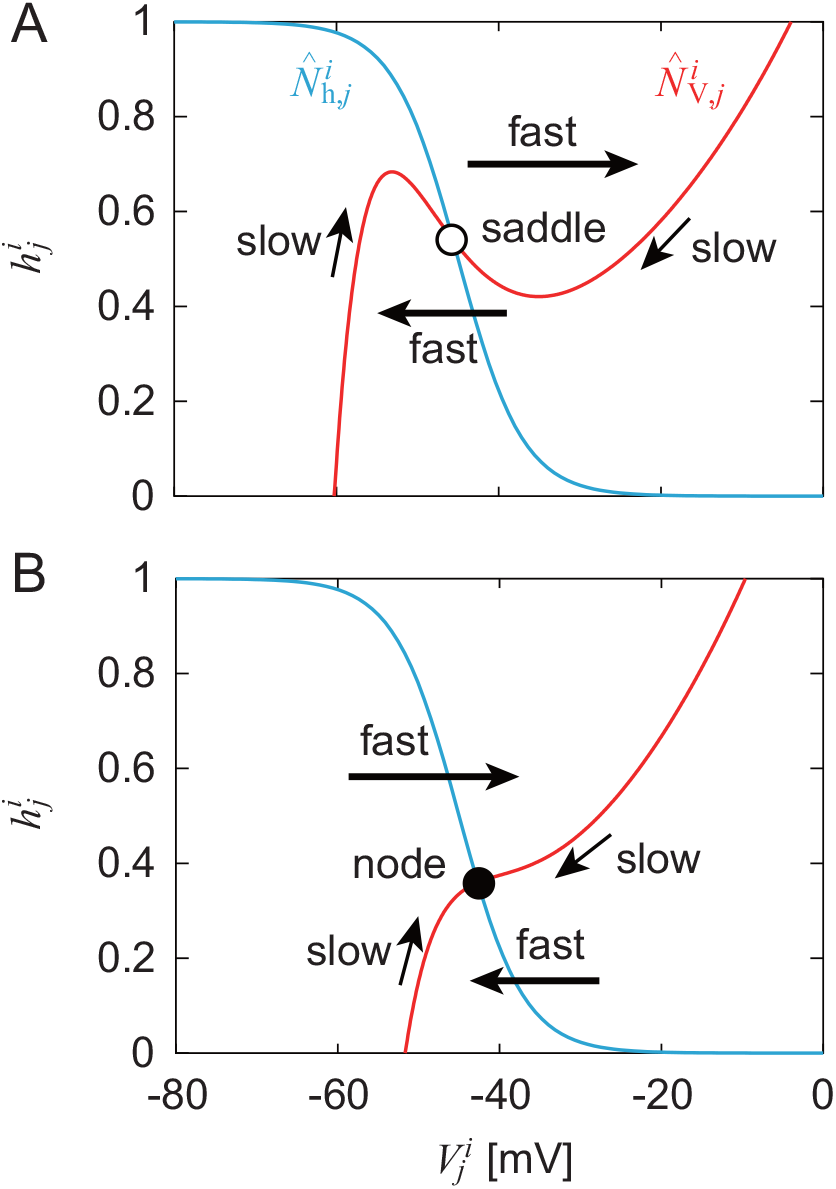
Nullclines 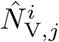 and 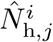 in 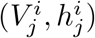 for (*i, j*) = (left, RG-F) and (right, RG-E). A. 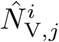 has two extremes and intersection with 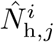 is saddle. B. 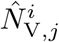changes monotonically and intersection with 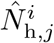is stable node. Bold and thin arrows represent fast and slow dynamics, respectively.

### 4.2. Adaptation mechanism based on nullcline analysis

Figure 7A shows the time profiles of the membrane potentials of the left RGF (ipsilateral, i.e., affected side) and right RG-E (contralateral side) and afferent feedback from the flexor and extensor muscles. Figures 7B and C show the trajectories of the state 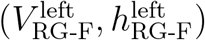and nullclines 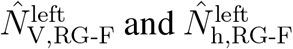 of the left RG-F, and the trajectories of the state 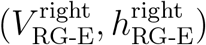 and nullclines 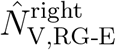 and 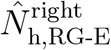 of the right RG-E, respectively. Although the trajectories show a closed loop during flat-surface walking due to a stable limit cycle, they are disturbed after a foot enters a hole (indicated by ‘a’).

**Figure 7.**
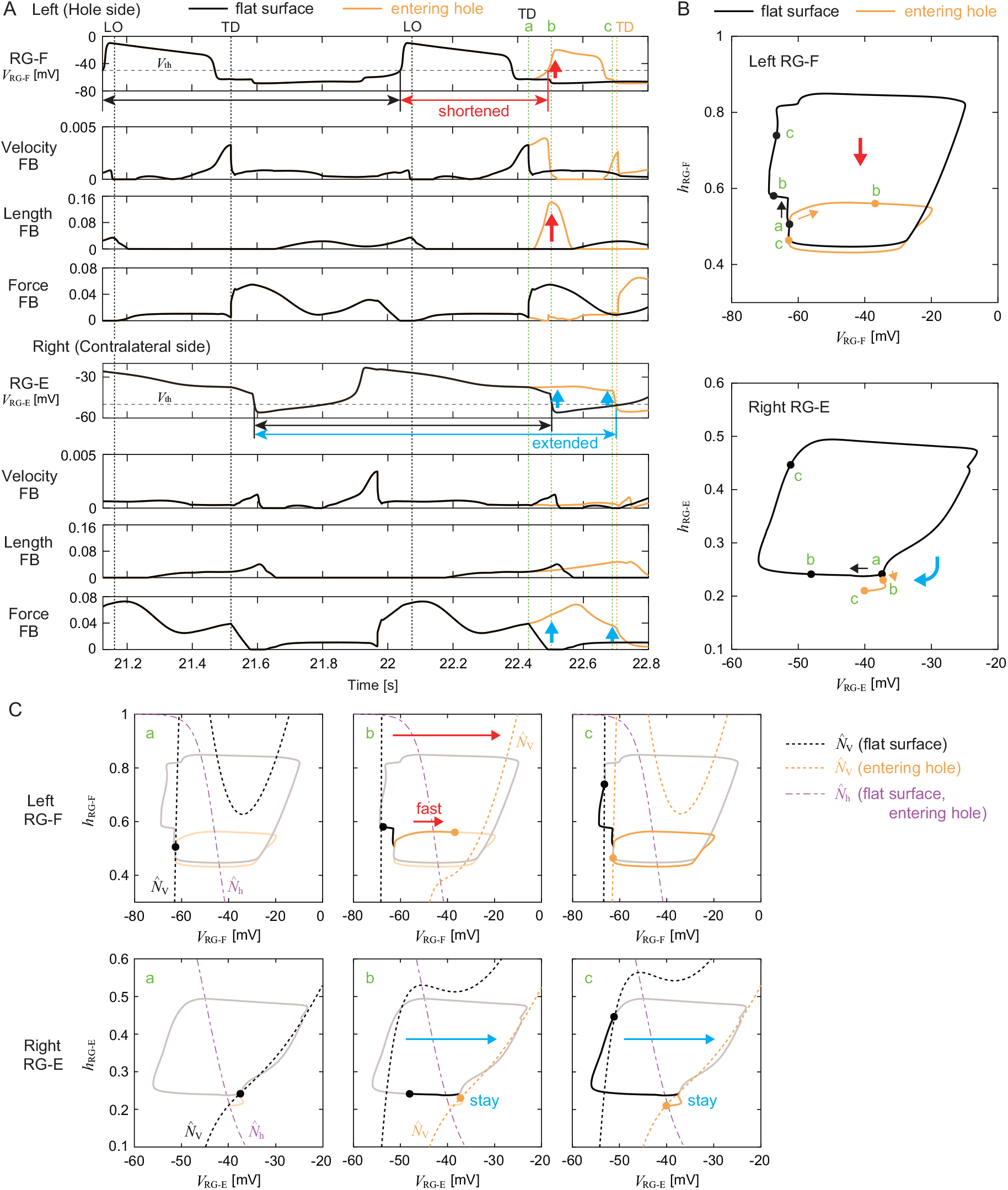
Nullcline analysis of CPG dynamics by comparing the response to the left foot entering a hole with the steady oscillation during flat-surface walking. A. Time profiles of RG membrane potentials and afferent feedback from flexor and extensor muscles. ‘a’, ‘b’, and ‘c’ indicate the times at which the left foot enters a hole (touchdown in flat-surface walking), at which the foot is lifted (early stance phase in flat-surface walking), and just before the foot touches the belt (midstance phase in flat-surface walking), respectively. TD and LO indicate the touchdown and liftoff of the left hindlimb, respectively. B. Trajectories of state 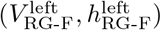 and nullclines 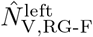 and 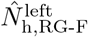 of left RG-F. C. Trajectories of state 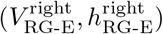 and nullclines 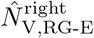 and 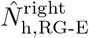 of right RG-E.

On the ipsilateral side that fell into the hole, the length feedback increased and the nullcline 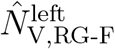 shifted to the right (indicated by ‘b’). This means that the hyperextension of the hindlimb was sensed by the length feedback of the flexor muscles which excited the flexor center and thereby shifted the nullcline. As a result, the state switched to fast dynamics, which rapidly increased 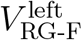. The trajectory shows a shortcut compared to the closed loop during flat-surface walking. These changes explain the early onset of flexor muscle activity in Fig. 4A, which allowed early hindlimb flexion to lift the foot out of the hole. These results are consistent with those of our previous work [28].

On the contralateral side, the force feedback increased and the nullcline 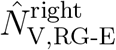 remained on the right side until the ipsilateral foot touched the belt (indicated by ‘b’ and ‘c’). This means that the loading was sensed by the force feedback of the extensor muscles which excited the extensor center and thereby maintained the nullcline. As a result, the state remained in slow dynamics, which maintained a large value of 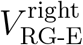. These changes explain the prolonged extensor muscle activity in Fig. 4B, which allowed the body weight to continue to be supported by the contralateral hindlimb.

We also performed a nullcline analysis on the models without length feedback from the flexor muscles or without force feedback from the extensor muscles that failed to recover from entering a hole in Figs. 5A and B (see details in Appendix D). In the model without length feedback, the nullcline 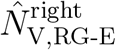 on the contralateral side remained on the right side and extensor muscle activity was prolonged (Fig. SD.10), as in Fig. 7. However, the nullcline 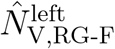 on the ipsilateral side did not shift to the right and early flexor activity did not occur, unlike in Fig. 7. This explains why the model without length feedback from the flexor muscles failed to recover.

In the model without force feedback, the nullcline 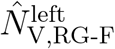 on the ipsilateral side shifted to the right due to an increase in length feedback from the flexor muscles, resulting in a shortcut in the trajectory and early flexor activity (Figs. 8A and SD.11), as in Fig. 7. In addition, despite the absence of force feedback, the nullcline 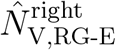 on the contralateral side remained on the right side and extensor muscle activity was prolonged (Figs. 8A and SD.11), as in Fig. 7. In fact, the maintenance of the nullcline and the prolongation of extensor activity on the contralateral side were caused by the early flexor activity on the ipsilateral side. Specifically, length feedback from the ipsilateral flexor muscles excited the ipsilateral flexor center, which in turn excited the contralateral extensor center via commissural interneuron (C1) (Fig. 8B). In other words, this reflects interlimb coordination via reciprocal movement, in which flexion of one hindlimb induces extension of the contralateral hindlimb. This reciprocal mechanism also contributed to the model with all feedback in Fig. 7. However, the prolongation of extensor activity on the contralateral side due to reciprocal coordination alone was insufficient to produce the level of muscle activity required to support body weight; additional contributions from force feedback to the contralateral PF and MN were also necessary (Fig. 8C).

**Figure 8.**
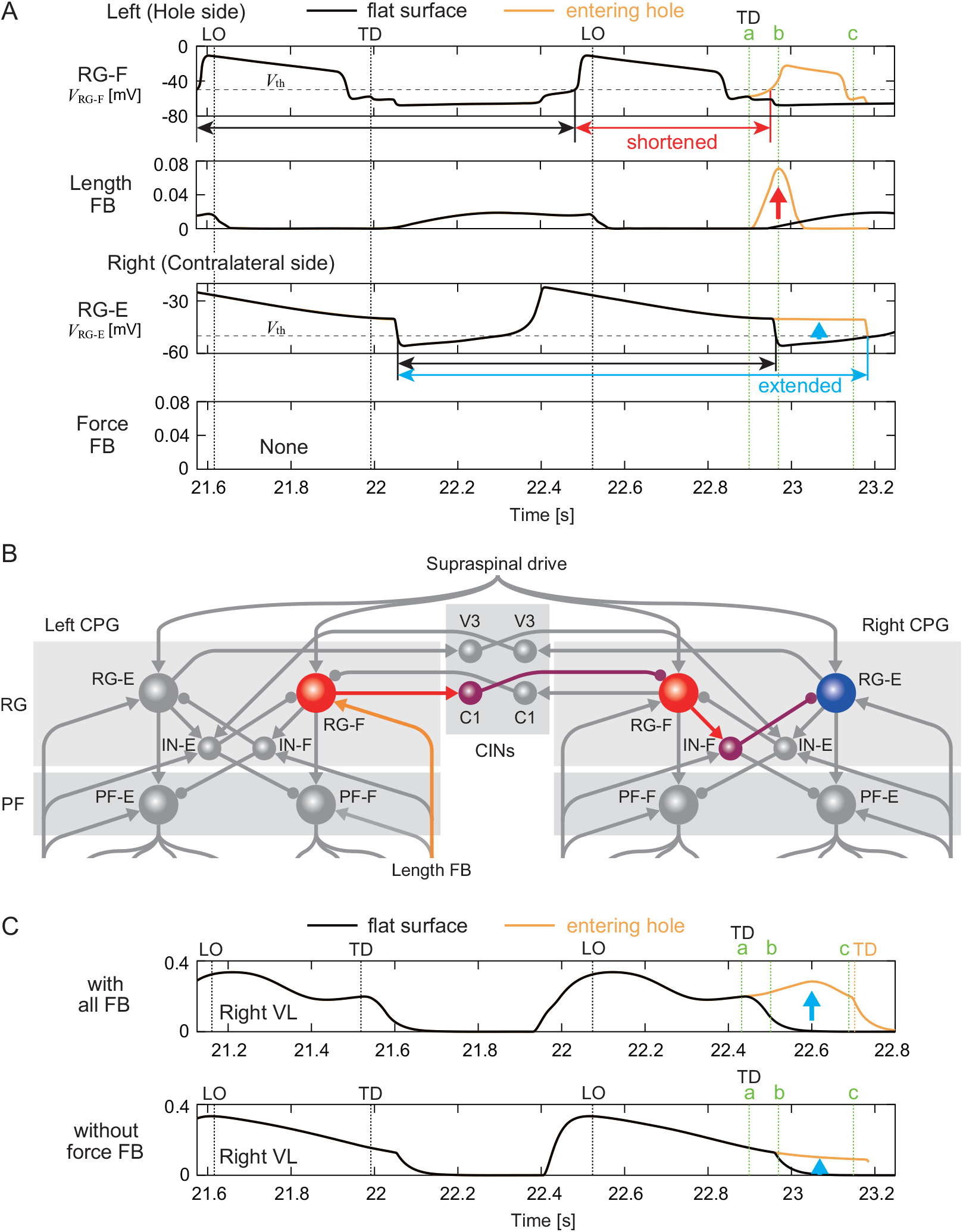
Reciprocal interlimb coordination during the response to the left foot entering a hole. A. Time profiles of RG membrane potentials and afferent feedback from flexor and extensor muscles of the model without force feedback. ‘a’, ‘b’, and ‘c’ indicate the times at which the left foot enters a hole (touchdown in flat-surface walking), at which the hindlimb is hyperextended (early stance phase in flat-surface walking), and at which the hip drops (midstance phase in flat-surface walking), respectively. TD and LO indicate the touchdown and liftoff of the left hindlimb, respectively. B. Reciprocal network in which length feedback from the ipsilateral flexor muscles excites the contralateral extensor center via commissural interneuron (C1). C. Muscle activity of the contralateral VL muscle compared between the model with all feedback and the model without force feedback.

## 5. Discussion

When a foot falls into a hole while walking on a treadmill, it must be quickly pulled out and returned to the treadmill belt to continue walking. Meanwhile, the contralateral hindlimb needs to support the body weight to prevent falling. To achieve such an adaptive response to this environmental disturbance, spinal CPGs should modify motor commands based on sensory feedback. Since the hindlimb that falls into the hole and loses ground contact becomes hyperextended, flexor activity must begin early. On the contralateral side, extensor activity must continue until the foot is pulled out of the hole. In cat locomotion, it has been suggested that sensing stretch of the flexor muscles via Ia and II sensory afferents [21, 26] and limb unloading via Ib sensory afferents of extensor muscles [12, 42, 60] play an important role in the transition from extensor activity during the stance phase to flexor activity during the swing phase. In our model, the velocityand lengthdependent (Ia, II) feedback of flexor muscles and the force-dependent (Ib) feedback of extensor muscles were involved in this phase transition and contributed to the adaptive response to the loss of ground support. Specifically, increasing the length feedback from the flexor muscles caused the flexor activity to begin just after the foot entered the hole (‘b’ in Fig. 7A). This enabled the foot to be quickly pulled out of the hole and placed back onto the treadmill belt. Simultaneously, increasing the force feedback from the extensor muscles on the contralateral side, along with the reciprocal contribution from the ipsilateral flexor center (Fig. 8) maintained the contralateral extensor activity (‘b’ and ‘c’ in Fig. 7A). This enabled the body weight to be supported until the foot was pulled out of the hole. In contrast, the model without these afferent feedbacks failed to continue walking once a foot stepped into a hole (Movies 6 and 7). These results are consistent with previous neurophysiological findings [12, 21, 26, 42, 60].

In general, stepping movements consist of flexion in the swing phase and extension in the stance phase that alternate between the left and right hindlimbs during walking. However, the adaptive response to the loss of ground support requires this alternating relationship to be broken temporarily. Specifically, while two flexions occur on the affected (hole) side before and after entering the hole, it is necessary to prolong extension on the contralateral side. In addition to simulation of the neuromusculoskeletal model, the nullcline analysis based on the dynamic systems theory revealed that, on the side of the hole, the change in the nullcline due to increased length feedback from the flexor muscles caused the RG neuron to switch to fast dynamics, resulting in a shortcut in the trajectory (Fig. 7C). This led to early flexor activity, i.e. two consecutive flexions occurred on the side of the hole. Furthermore, it was revealed that, on the contralateral side, increased force feedback from the extensor muscles, along with the reciprocal contribution from the ipsilateral flexor center (Fig. 8), maintained the nullcline, enabling the RG neuron to remain in slow dynamics (Fig. 7C). This allowed the extensor activity to continue. By controlling transitions between fast and slow CPG dynamics through sensory feedback, the system selectively disrupted leftright alternation in a context-dependent manner, an insight that would be difficult to extract without this dynamical systems perspective. This active disruption of the left-right alternation has also been observed during split-belt treadmill walking in cats [15, 16, 17], suggesting that this is an important function for adaptive locomotion.

The two-level half-center CPG model used in this study has played a key role in previous works [3, 5, 6, 7, 32, 38, 50, 51, 52, 64] and helped extend our understanding of the neural networks involved in interlimb coordination in mammalian gait generation. However, these studies did not incorporate a mechanical model of the body, ignoring the mechanisms of interlimb coordination based on interactions between the nervous system, the musculoskeletal system, and the environment. In the present study, we incorporated a musculoskeletal model and employed afferent feedback to gain information on body dynamics. This allowed us to understand and suggest the role of these interactions. The afferent feedback was projected exclusively to the ipsilateral CPG and not directly to the contralateral CPG. Yet, coordinated interlimb responses still emerged, suggesting that physical coupling of the hindlimbs and commissural connections were sufficient to transmit information across sides. In this way, adaptive responses to the loss of ground support were achieved through a combination of local information processing in each hindlimb and global system dynamics mediated by body movements. These findings were made possible for the first time through the incorporation of the musculoskeletal model.

So far, various neuromechanical models incorporating a musculoskeletal model have been used to understand the mechanisms of adaptive locomotion through sensory feedback in mammals [2, 1, 8, 9, 13, 19, 27, 36, 39, 45, 53, 56, 61]. However, these neural models have been limited to abstract representations using simplified reflex loops or neural oscillators, which limited comparisons with real experimental data [4]. Unlike previous models, our model integrated a two-level physiologically-inspired CPG with structured afferent feedback from a biomechanically realistic musculoskeletal system. This architecture enabled us to mechanistically explain how the loss of ground support on one side drives an early flexor transition, while extensor activity is preserved on the contralateral side, not through hardwired transitions, but via dynamic modulation of the CPGs’ operating regime. This provides a conceptual advancement over prior neuromechanical studies, which often could not account for such transient asymmetric behaviors with high biological plausibility. A particularly notable feature is that our model was optimized using a randomized stepping surface but not explicitly trained on the hole condition. Nevertheless, it successfully generated adaptive interlimb responses when encountering a sudden loss of ground support. This generalization suggests that the modeled afferent feedback architecture, together with the intrinsic properties of the CPG, can support a repertoire of adaptive behaviors without context-specific tuning. Such robustness is a hallmark of biological systems and highlights the capacity of spinal circuits to reorganize their dynamics through local sensory signals.

Although we clarified the mechanism underlying the adaptive interlimb response to the sudden loss of ground support, our model has limitations. For example, it is a two-dimensional model constrained to the sagittal plane, whereas adaptive locomotion also requires lateral balancing [41]. Moreover, although our model used a simple PF model that generates basic alternating activity between flexors and extensors, actual muscle activity exhibits more complex patterns [10, 11, 29, 35] that could facilitate adaptive motor control. In addition, the simulation results of joint angles and muscle activity do not perfectly match experimental data (Figs. 3 and 4). In particular, the simulation results show greater vertical hip movement when the foot falls into the hole than the measured data. This discrepancy is largely due to the simplification of our model, in which the forelimbs are fixed to the trunk. This causes the model to rotate around the tips of the forelimbs, which exaggerates the vertical movement of the hip. Furthermore, while length and force feedback played important roles in the adaptive response to the loss of ground support and in flat-surface walking, velocity feedback contributed little to this study. These limitations suggest directions for future improvement, including full-body 3D mechanics [57, 58], more detailed PF circuitry, and context-sensitive feedback integration.

Our results also generate testable predictions. The nullcline-based mechanism implies that afferent-driven changes in network dynamics, and not just input amplitude, govern adaptive transitions. In this context, experimental strategies that selectively block length feedback are expected to delay or impede the initiation of flexor activity, while disruption of contralateral Ib input is likely to compromise weight support.

In this study, we focused on adaptive response to the loss of ground support during treadmill cat locomotion. However, our model and analysis could also be applied to other environmental changes and perturbations. In particular, the hole used in this study induced left-right asymmetric movements, requiring appropriate interlimb coordination. Therefore, our model could also contribute to our understanding of the adaptation mechanism in walking under asymmetric conditions, such as split-belt treadmill walking. Although we used a model with only hindlimbs and fixed forelimbs, we have already developed four-limb musculoskeletal models that incorporate both forelimbs and hindlimbs [57, 58, 59]. Incorporating the two-level half-center CPGs used in this study into the four-limb models will allow us to examine the formation and transition mechanisms of various gaits, such as walking, trotting, and galloping, through interlimb coordination across all four limbs. More broadly, our findings support a view of spinal locomotor circuits as flexible dynamical substrates shaped by continuous sensory input, rather than rigid pattern generators. Adaptive control emerges not from preprogrammed switches, but from feedback-induced transitions between dynamical regimes. This principle may underlie robustness in both natural and engineered locomotion systems.

## Appendix A. Parameters for the motor control model

Based on [5, 6, 20, 28, 34, 38], we determined the parameters for the synaptic connections 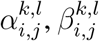, and *γ*_*j*_ (*j* ∈ *{*RG*}, {*IN*}, {*CIN*}, {*PF*}, l* ∈ *{*RG*}, {*IN*}, {*CIN*}*) and those for the currents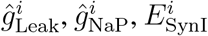, and 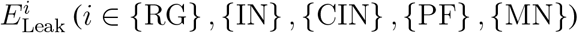 (*i* ∈ *{*RG*}, {*IN*}, {*CIN*}, {*PF*}, {*MN*}*) as shown in Tables A.1 and A.2, respectively. We used the following parameters:

**Table A.1.** Parameters for synaptic connections.

| Source neuron $l$ | Target neuron $j$ | | | | | | | |
| --- | --- | --- | --- | --- | --- | --- | --- | --- |
|  | RG-F | RG-E | IN-F | IN-E | C1 | V3 | PF-F | PF-E |
| Excitatory connection $\alpha_{i,j}^{k,l}$ | | | | | | | | |
| RG-F | - | - | 0.4 | - | 1 | - | 0.7 | - |
| RG-E | - | - | - | 0.4 | - | 0.5 | 0.7 | - |
| Contralateral V3 | - | - | - | 0.15 | - | - | - | - |
| Inhibitory connection $\beta_{i,j}^{k,l}$ | | | | | | | | |
| IN-F | - | 0.7 | - | - | - | - | - | 2.1 |
| IN-E | 0.6 | - | - | - | - | - | 0.3 | - |
| Contralateral C1 | 1 | - | - | - | - | - | - | - |
| Supraspinal connection $\gamma_j$ | 0.02 | 0.15 | - | - | - | - | - | - |

**Table A.2.** Parameters for currents.

| | Neuron $i$ | | | | |
| --- | --- | --- | --- | --- | --- |
| | $i \in \{\text{RG}\}$ | $i \in \{\text{IN}\}$ | $i \in \{\text{CIN}\}$ | $i \in \{\text{PF}\}$ | $i \in \{\text{MN}\}$ |
| $\hat{g}_{\text{Leak}}^i$ | 4.5 | 2.8 | 2.8 | 1.6 | 1.6 |
| $\hat{g}_{\text{NaP}}^i$ | 4.5 | - | - | 0.5 | 0.3 |
| $E_{\text{SynI}}^i$ | -75 | -75 | -75 | -70 | -70 |
| $E_{\text{Leak}}^i$ | -62.5 | -60 | -60 | -64 | -64 |

## Appendix B Parameters determined by optimization

Table B.3 shows the excitatory connection 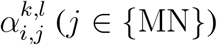 for the ipsilateral side (*i* = *k*) (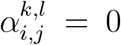 for the contralateral side (*i ≠ k*)) and the parameters for the afferent feedback 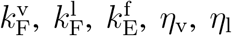, and *η*_f_, which were determined by optimization.

**Table B.3.** CPG Parameters determined by optimization.

| $\alpha_{i,j}^{k,l}$<br>Source neuron $l$ | Target neuron $j$ | | | | | | |
| --- | --- | --- | --- | --- | --- | --- | --- |
|  | MN-IP | MN-GM | MN-VL | MN-TA | MN-SO | MN-BF | MN-GA |
| PF-F | 0.09 | - | - | 0.10 | - | 0.08 | - |
| PF-E | - | 0.07 | 0.12 | - | 0.08 | - | 0.12 |
| | $k_{\text{F}}^{\text{v}}$ | $k_{\text{F}}^{\text{l}}$ | $k_{\text{E}}^{\text{f}}$ | $\eta_{\text{v}}$ | $\eta_{\text{l}}$ | $\eta_{\text{f}}$ | |
|  | 0.0007 | 0.53 | 0.16 | 1.0 | 1.1 | 1.1 |  |

## Appendix C Reproduction of flat-surface treadmill walking without afferent feedback

We determined the parameters of the model without velocity feedback, the model without length feedback, and the model without force feedback by optimization with the walking environment (Fig. 2A), where the belt level changes in each step, in the same way as the model with all afferent feedback. Table C.4 shows the parameters of each model determined by optimization.

**Table C.4.** CPG parameters of the models without afferent feedback determined by optimization.

| $\alpha_{i,j}^{k,l}$<br>Source neuron $l$ | Target neuron $j$ | | | | | | |
| --- | --- | --- | --- | --- | --- | --- | --- |
|  | MN-IP | MN-GM | MN-VL | MN-TA | MN-SO | MN-BF | MN-GA |
| without velocity FB |  |  |  |  |  |  |  |
| PF-F | 0.10 | - | - | 0.09 | - | 0.08 | - |
| PF-E | - | 0.07 | 0.12 | - | 0.08 | - | 0.12 |
| without length FB |  |  |  |  |  |  |  |
| PF-F | 0.09 | - | - | 0.09 | - | 0.07 | - |
| PF-E | - | 0.08 | 0.13 | - | 0.08 | - | 0.12 |
| without force FB |  |  |  |  |  |  |  |
| PF-F | 0.10 | - | - | 0.09 | - | 0.08 | - |
| PF-E | - | 0.07 | 0.18 | - | 0.10 | - | 0.16 |
| | $k_F^v$ | $k_F^l$ | $k_E^f$ | $\eta_v$ | $\eta_l$ | $\eta_f$ | |
| without velocity FB | - | 0.45 | 0.16 | - | 1.2 | 1.0 |  |
| without length FB | 0.026 | - | 0.10 | 1.3 | - | 1.3 |  |
| without force FB | 0.012 | 0.28 | - | 1.7 | 1.4 | - |  |

Figure C.9 shows the simulation results of flat-surface treadmill walking for each model without changing the belt level, compared with the results for the model with all afferent feedback in Fig. 3. Even without any afferent feedback, flat-surface walking was almost the same as with all afferent feedback.

**Figure C.9.**
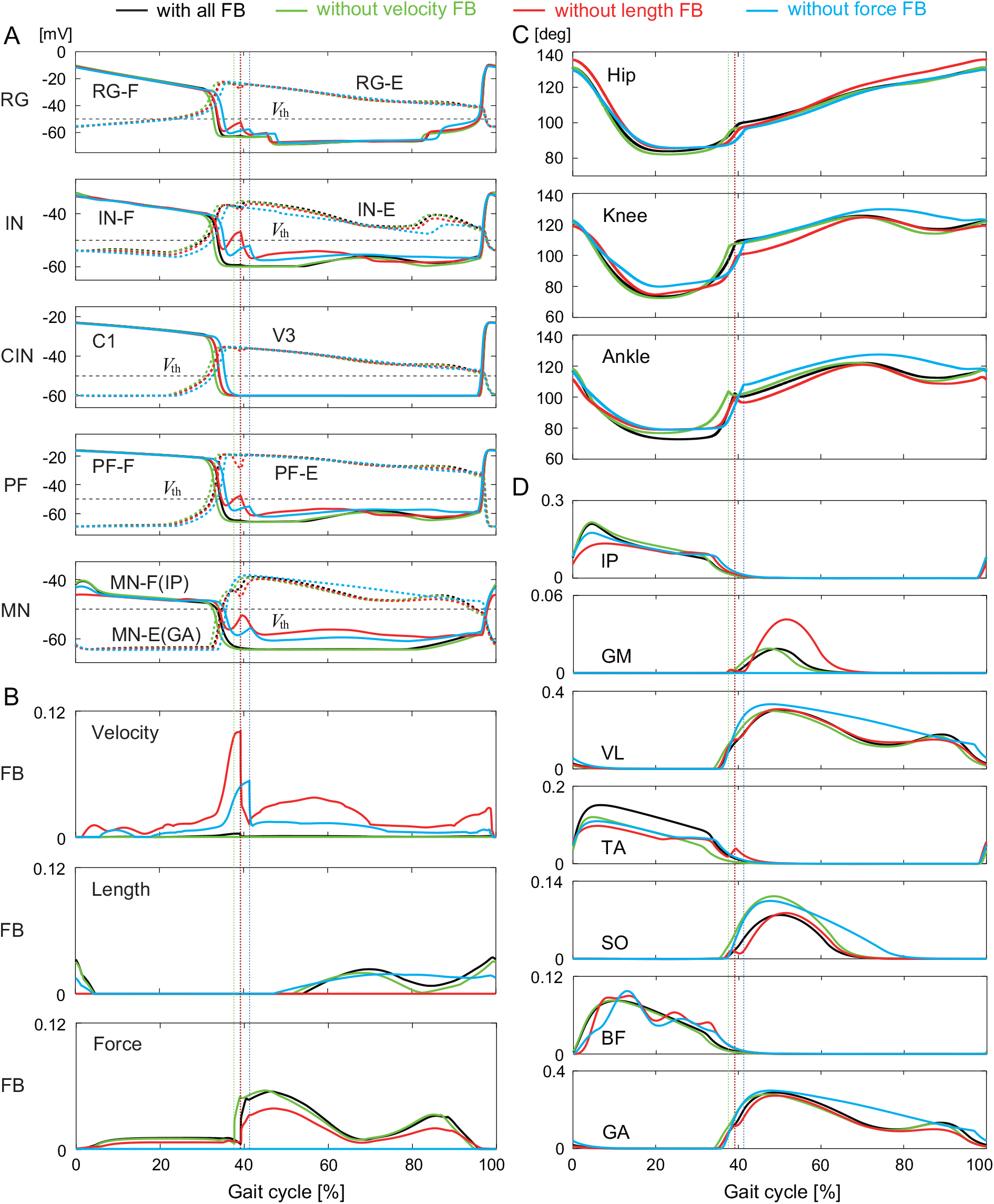
Simulation results for flat-surface walking of the models without afferent feedback compared with those of the model with all afferent feedback. A. Membrane potentials of CPG neurons and B. afferent feedback from flexor and extensor muscles. C. Joint angles and D. muscle activities. Liftoffs are represented by 0% and 100% in the gait cycle. Vertical lines indicate touchdowns.

## APPENDIX D Nullcline analysis on the model without afferent feedback

We performed a nullcline analysis on the models without length feedback from the flexor muscles or without force feedback from the extensor muscles, in the same way as the analysis in Fig. 7 for the model with all afferent feedback. Specifically, we investigated the dynamics of the CPG models by focusing on the the nullclines of the left RG-F and the right RG-E using the simulation results during the response in Fig. 5 and compared them with those during flat-surface walking in Fig. C.9.

Figure D.10A shows the time profiles of the membrane potentials of the left RG-F (ipsilateral, i.e., affected side) and right RG-E (contralateral side) and afferent feedback from the flexor and extensor muscles, for the model without length feedback. Figures D.10B and C show the trajectories of the state 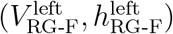 and nullclines 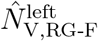 and 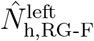 of the left RG-F, and the trajectories of the state 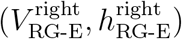 and nullclines 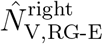 and 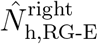 of the right RG-E, respectively, for the model without length feedback. Figures D.11A, B, and C show the same figures for the model without force feedback. Although the trajectories show a closed loop during flat-surface walking due to a stable limit cycle, they are disturbed after a foot enters a hole (indicated by ‘a’).

**Figure D.10.**
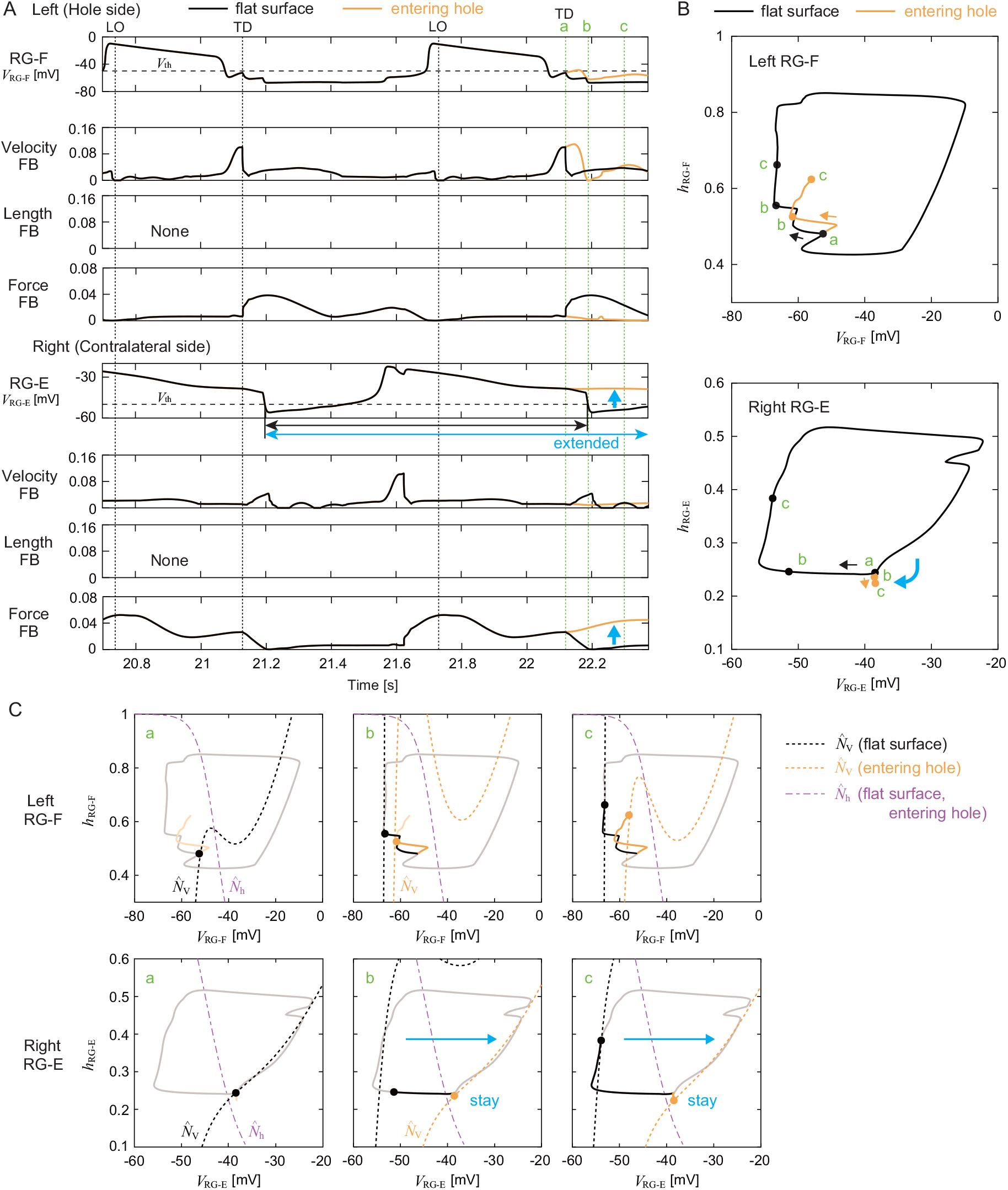
Nullcline analysis of CPG dynamics for the model without length feedback of the flexor muscles by comparing the response to left foot entering a hole with the steady oscillation during flat-surface walking. A. Time profiles of RG membrane potentials and afferent feedback from flexor and extensor muscles. ‘a’, ‘b’, and ‘c’ indicate the times at which left foot enters a hole (touchdown in flat-surface walking), at which the hindlimb is hyperextended (early stance phase in flat-surface walking), and at which the hip drops (midstance phase in flat-surface walking), respectively. TD and LO indicate the touchdown and liftoff of the left hindlimb, respectively. B. Trajectories of state 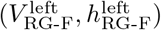 and nullclines 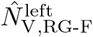 and 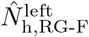 of left RG-F. C. Trajectories of state 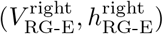 and nullclines 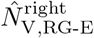 and 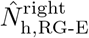 of right RG-E.

**Figure D.11.**
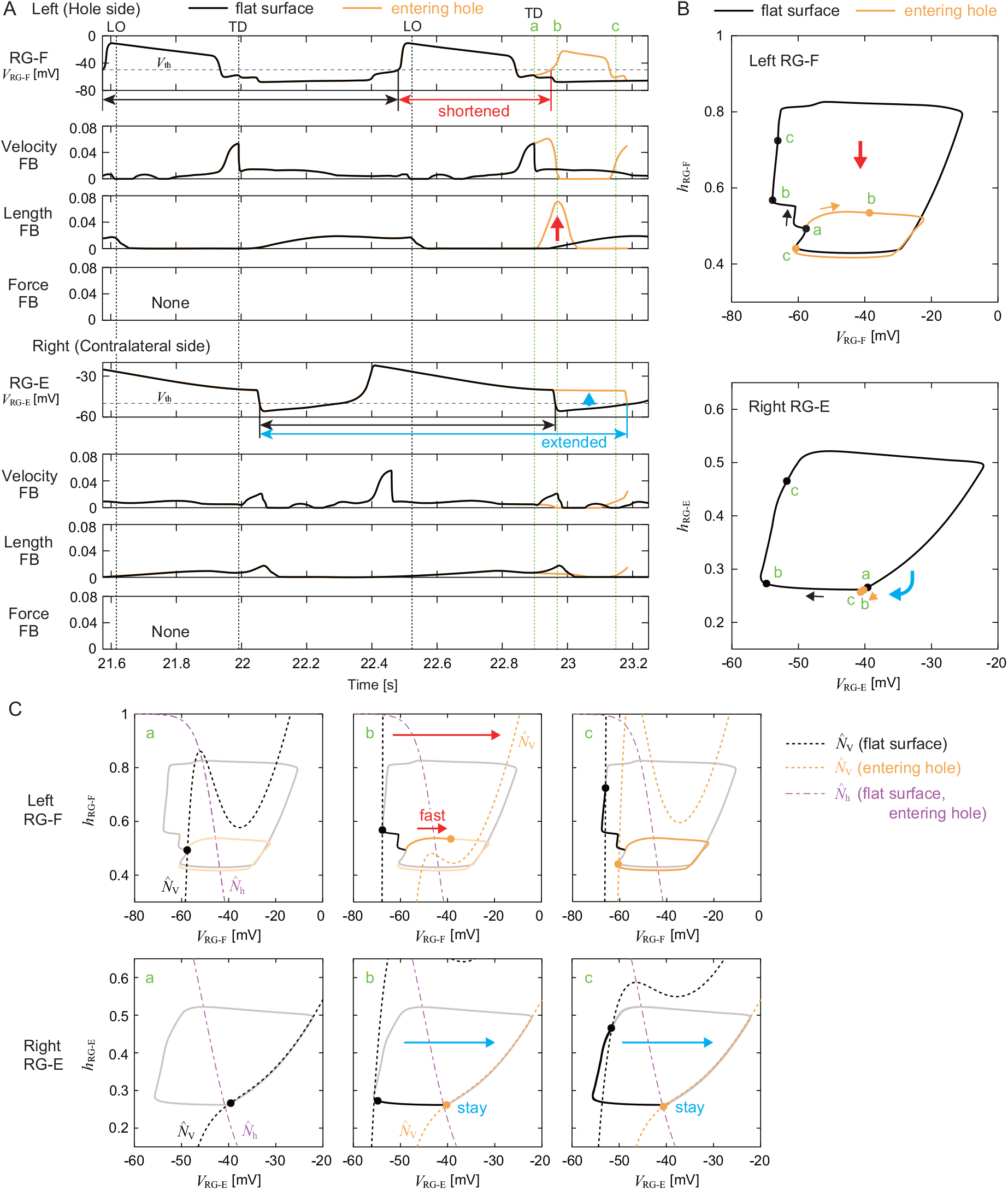
Nullcline analysis of CPG dynamics for the model without force feedback of the extensor muscles by comparing the response to left foot entering a hole with the steady oscillation during flat-surface walking. A. Time profiles of RG membrane potentials and afferent feedback from flexor and extensor muscles. ‘a’, ‘b’, and ‘c’ indicate the times at which left foot enters a hole (touchdown in flat-surface walking), at which the hindlimb is hyperextended (early stance phase in flat-surface walking), and at which the hip drops (midstance phase in flat-surface walking), respectively. TD and LO indicate the touchdown and liftoff of the left hindlimb, respectively. B. Trajectories of state 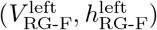 and nullclines 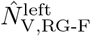 and 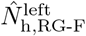 of left RG-F. C. Trajectories of state 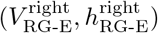 and nullclines 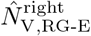 and 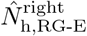 of right RG-E.

When the model did not receive length feedback of the flexor muscles, the nullcline 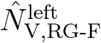on the ipsilateral side that fell into the hole did not shift to the right, unlike in Fig. 7. As a result, the state remained in slow dynamics (‘b’ and ‘c’) and early flexor activity did not occur. On the contralateral side, the force feedback of the extensor muscles increased and the nullcline 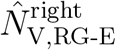 remained on the right side (‘b’ and ‘c’), as in Fig. 7. As a result, the state remained in slow dynamics, which maintained a large value of 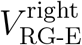 and prolonged extensor muscle activity.

For the model without force feedback of the extensor muscles, the length feedback increased on the ipsilateral side and the nullcline 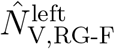 shifted to the right (‘b’), as in Fig. 7. As a result, the state switched to fast dynamics, which rapidly increased 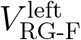. The trajectory showed a shortcut, resulting in early flexor activity. On the contralateral side, the nullcline 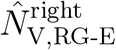 remained on the right side as in Fig. 7, despite the absence of force feedback (‘b’ and ‘c’). As a result, the state remained in slow dynamics, which maintained a large value of 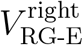 and prolonged extensor muscle activity.

## Acknowledgments

This study was supported in part by JSPS KAKENHI Grant Numbers JP24H00297 and JP19KK0377 and JST FOREST Program Grant Number JPMJFR2021 to SA, NIH Grant Numbers R01NS112304 and R01NS115900 to SMD, R01NS130799 and R01NS110550 to IAR, and T32NS121768 to ABL, and NSF CRCNS/DARE Grant Number 2113069 to JA and IAR.

## Data availability

Data will be made available on request.

## Author contributions

SA developed the study design. KS, YA, YK, and SA developed the cat neuro-musculoskeletal model in consultation with STR, ABL, SNM, JA, IAR, and SMD. KS, YA, YK, and FM performed computer simulation and analyzed the data. KS and SA wrote the manuscript and all the authors reviewed, edited, and approved it.

